# Kif9 is an active kinesin motor required for ciliary beating and proximodistal patterning of motile axonemes

**DOI:** 10.1101/2021.08.26.457815

**Authors:** Mia J. Konjikusic, Chanjae Lee, Yang Yue, Bikram D. Shrestha, Ange M. Nguimtsop, Amjad Horani, Steven Brody, Vivek N. Prakash, Ryan S. Gray, Kristen J. Verhey, John B. Wallingford

## Abstract

Most motile cilia have a stereotyped structure of nine microtubule outer doublets and a single central pair of microtubules. The central pair microtubules are surrounded by a set of proteins, termed the central pair apparatus. A specific kinesin, Klp1 projects from the central pair and contributes to ciliary motility in *Chlamydomonas*. The vertebrate orthologue, Kif9 is required for beating in mouse sperm flagella, but the mechanism of Kif9/Klp1 function remains poorly defined. Here, using *Xenopus* epidermal multiciliated cells, we show that Kif9 is necessary for ciliary motility as well as leads to defects in the distal localization of not only central pair proteins, but also radial spokes and dynein arms. In addition, single-molecule assays *in vitro* revealed that *Xenopus* Kif9 is a processive motor, though like axonemal dyneins it displays no processivity in ciliary axonemes *in vivo*. Thus, our data suggest that Kif9 plays both indirect and direct role in ciliary motility.

## Introduction

Motile cilia are essential for fluid flow across epithelia and for cell motility in a variety of contexts in the animal kingdom (Brooks and Wallingford, 2014; Spassky and Meunier, 2017; Ishikawa, 2017). Most motile cilia have a conserved radial structure of nine microtubule outer doublets and a single central pair of microtubules. Ciliary motility relies on several different protein complexes that are organized throughout the axoneme in a specific fashion that are crucial for proper motility. These include the inner and outer dynein arms on the outer doublets and radial spokes that connect to inner dynein arms and project towards to central pair (Ishikawa, 2017). In addition, several proteins surround the central pair to form the central pair apparatus (CA) (Loreng and Smith, 2017). All of these elements work in concert to orchestrate proper ciliary beat frequency and waveform.

In addition to the radial patterning of motile axonemes, a complex proximal to distal patterning has been extensively studied in *Chlamydomonas* and *Tetrahymena* (Pedersen et al., 2003; Bui et al., 2012; Louka et al., 2018). Though less well-defined, such proximodistal patterning is also critical in vertebrate multiciliated cells (Fliegauf et al., 2005; Loges et al., 2008; Yamamoto et al., 2010). The function and composition of the distal tip of motile cilia is of particular interest, as it is thought to be critical to protect the axoneme from disintegration due to the massive forces associated with beating. The distal tip differs in structure wildly across ciliary types (Soares et al., 2019), but enrichment of the microtubule end-binding proteins Eb1 and Eb3 in distal regions is conserved from algae to vertebrates (Brooks and Wallingford, 2012; Schrøder et al., 2011; Pedersen et al., 2003). Finally, we and others have identified Spef1, a microtubule bundling protein, is also highly enriched in the distal tip of motile axonemes in *Xenopus* (Gray et al., 2009; Schrøder et al., 2011). Interestingly, Spef1 also contributes to central pair formation in mouse multiciliated cells. Whether Spef1 and the central apparatus contribute to tip integrity in multiciliated cells remains unclear.

Here, we examined the function of Kif9, the vertebrate homolog of the *Chlamydomonas* kinesin Klp1, which projects from the C2 microtubule of the central pair and is required for ciliary beating (Bernstein et al., 1994; Miyata et al., 2020; Yokoyama et al., 2004; Lechtreck and Witman, 2007). A recent report describes improper waveforms and disrupted ciliary beat in sperm from Kif9 mutant mice (Miyata et al., 2020), and as in *Chlamydomonas*, Kif9 mutant sperm do not display overt defects in the assembly of the central pair microtubules (Yokoyama et al., 2004; Miyata et al., 2020). Thus, the mechanisms by which loss of Klp1/Kif9 disrupts cilia beating remain unclear. Here, we find that loss of Kif9 in *Xenopus* results in defects in ciliary beating and cilia length related to specific disruption of the distal tip of the axoneme. Strikingly, we observed loss not only of central pair proteins, but also of radial spokes and dynein arms from distal axonemes after Kif9 knockdown. Finally, we show that Kif9 is a slowly processive microtubule motor *in vitro* but does not display processivity in the axoneme. These data provide new insights into ciliary beating and so may contribute to our understanding of human motile ciliopathies.

## Results

### Kif9 localizes to the axonemes of multiciliated cells in *Xenopus* and human airways

*Xenopus* tadpoles are a powerful model for studying the formation and function of multiciliated cells (Walentek and Quigley, 2017), so we used this system to examine the localization of Kif9 in multiciliated cells. *En-face* confocal imaging of Kif9-GFP mRNA-injected, whole mounted embryos revealed localization to cilia and basal bodies of multiciliated cells (Figure 1A-B’’). We confirmed endogenous Kif9 localization in ciliary axonemes using immunofluorescence and co-staining with acetylated tubulin (Figure 1C-C”). This localization is also evolutionarily conserved, as Kif9 immunostaining of human airway MCCs also revealed localization to cilia (Figure 1F-F”). Thus, Kif9 is expressed in *Xenopus* and human airway multiciliated cells where it localizes to ciliary axonemes.

**Figure 1.**
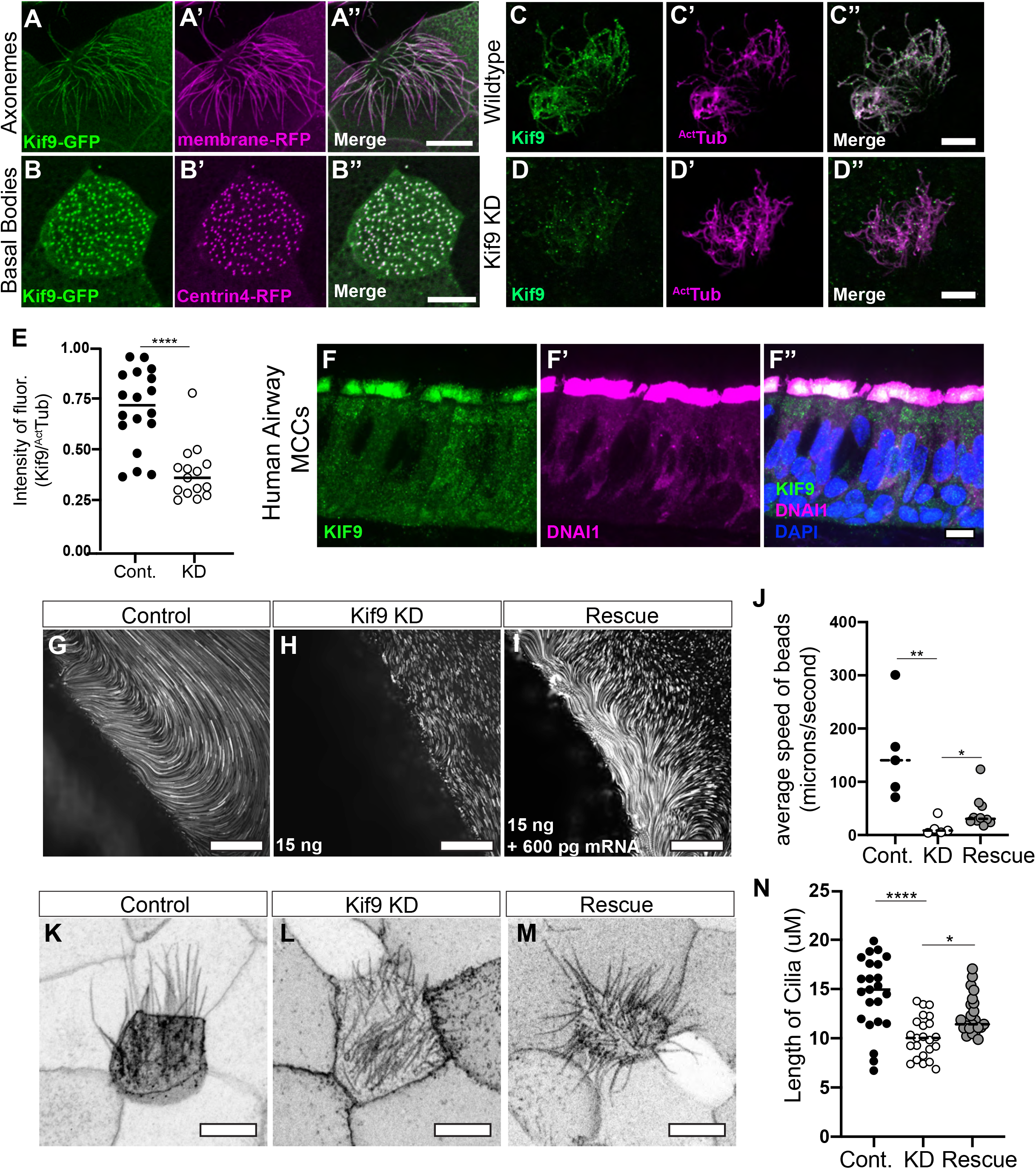
Kif9 localization to motile cilia and is essential for ciliary beating. A-A’’’. Kif9-GFP construct localizes to xenopus multiciliated cells in the axonemes. Kif9-GFP in green (A), membrane-RFP in magenta (A’). B-B’’. Kif9-GFP construct localizes to basal bodies of multiciliated cells in *Xenopus*. Kif9-GFP in Green (B), Centrin4-RFP in magenta (B’). Scale bars for A-B” 10uM. C-C’’. Kif9 antibody (green) confirms localization to xenopus axonemes. Kif9 in green (C) and acetylated tubulin in magenta (C’) Scale bars 10uM. D-D’’. Knockdown with 15 ng of morpholino reduces Kif9 (green) from cilium. E. Quantification of Kif9 staining intensity over the intensity of acetylated tubulin in control and morpholino injected frogs. F-F’’. KIF9 immunostaining in human airway multiciliated cells. F. KIF9 immunostaining (green). F’ DNAI immunostaining (magenta). F’’ Merge of F-F’ with DAPI. Scale Bar 10 microns. G-I. Still images from bead flow movies subjected to Flowtrace analysis. G. Control injected, H. 15 ng Kif9 morpholino injected, and I. 15ng morpholino with 600pg Kif9 mRNA (Rescue). Scale bars for G-I 50 microns. J. Graph of bead velocity averages from collected movies. P-values: ** = 0.0094 and * = 0.0280 K-M. Representative images of ciliary length marked with membrane-GFP in K. Control, L, Kif9 morpholino, and M. Rescue injected embryos. Scale bars 10 microns. N. Graph of ciliary length from Control, Morpholino, or Rescue cilia. P-values: ****< 0.0001 and * = 0.0174.

### Kif9 is necessary for ciliary beating in multiciliated cells

Multiciliated cells generate fluid flow across several different epithelia (Brooks and Wallingford, 2014; Spassky and Meunier, 2017) and the *Xenopus* epidermis offers an accessible model for studying those flows (Walentek and Quigley, 2017). In order to knockdown (KD) expression of Kif9 and study effects of its loss on cilia-mediated fluid flow, we designed a splice-blocking morpholino oligonucleotides against the first exon of Kif9 in *Xenopus*. Kif9 mRNA knockdown was confirmed by RT-PCR (Supp. Figure 1A). Crucially, imaging of Kif9 immunostaining in knockdown cilia also revealed reduced levels of Kif9 protein in cilia (Figure 1D-D”, 1E).

To visualize flows, we whole mounted embryos with beads mixed into the media and imaged at high speeds with confocal microscopy. Using Flowtrace analysis (Gilpin et al., 2017), we highlighted tracks created by beads across frames (Movie 1, 2 and 3). We then used Particle Image Velocimetry (PIV) analysis to quantify the ciliary beating-induced fluid flow speeds. In control embryos, we observed average fluid flow speeds of about 154 microns/second, whereas morpholino injected embryos showed a ten-fold reduction in flow (P-value = 0.0094; Figure 1G-H, 1J, Supp. Figure 1B). Reintroduction of Kif9-GFP mRNA into knockdown embryos rescued the flow defects, restoring flow rates to ∼42 microns/second, thereby supporting the specificity of this phenotype (Figure 1I, 1J, P-value = 0.0280).

To ask how Kif9 knockdown affects ciliary beating, we used high-speed imaging of cilia labeled with membrane-GFP. Compared to controls, Kif9 KD embryos displayed beat defects that were restored in the rescued embryos (Movies 4, 5, 6). These beat defects ranged from complete paralysis of cilia, to reduced beat frequency and disrupted waveforms. Kymographs of ciliary beating highlight the deficiencies in Kif9 knockdown embryos compared to controls. Beat patterns were restored in rescued embryos (Supp. Figure 1C).

Finally, in addition to ciliary beat defects, Kif9 KD embryos also had shorter cilia than controls (P-value < 0.0001, Figure 1K-L and 1N). This phenotype was also rescued by expression of Kif9-GFP (P-value = 0.0174; Figure 1M and 1N). These findings are consistent with the finding that axoneme length is often affected in central pair mutants in *Chlamydomonas* (Lechtreck et al., 2013). Together, these data demonstrate that Kif9 is necessary for normal ciliary length and beating in vertebrate multiciliated cells.

### Kif9 is necessary for central pair protein localization along the proximal/distal axis of the axoneme

We wished to assess how loss of Kif9 affects other components of the central pair. To this end, we generated GFP fusions of proteins known to act in specific sub-domains of the central apparatus and asked if their localization was altered by Kif9 KD. We tested the C2 bridge component Spag16 (Pf20 in *Chlamydomonas*) (Figure 2A). Live confocal imaging of embryos injected with membrane-RFP (gray) and GFP-Spag16 (orange) revealed a striking reduction in localization of Spag16, but intriguingly, this defect was restricted to the distal ends of the cilia (Figure 2B-E Supp. Figure 2).

**Figure 2.**
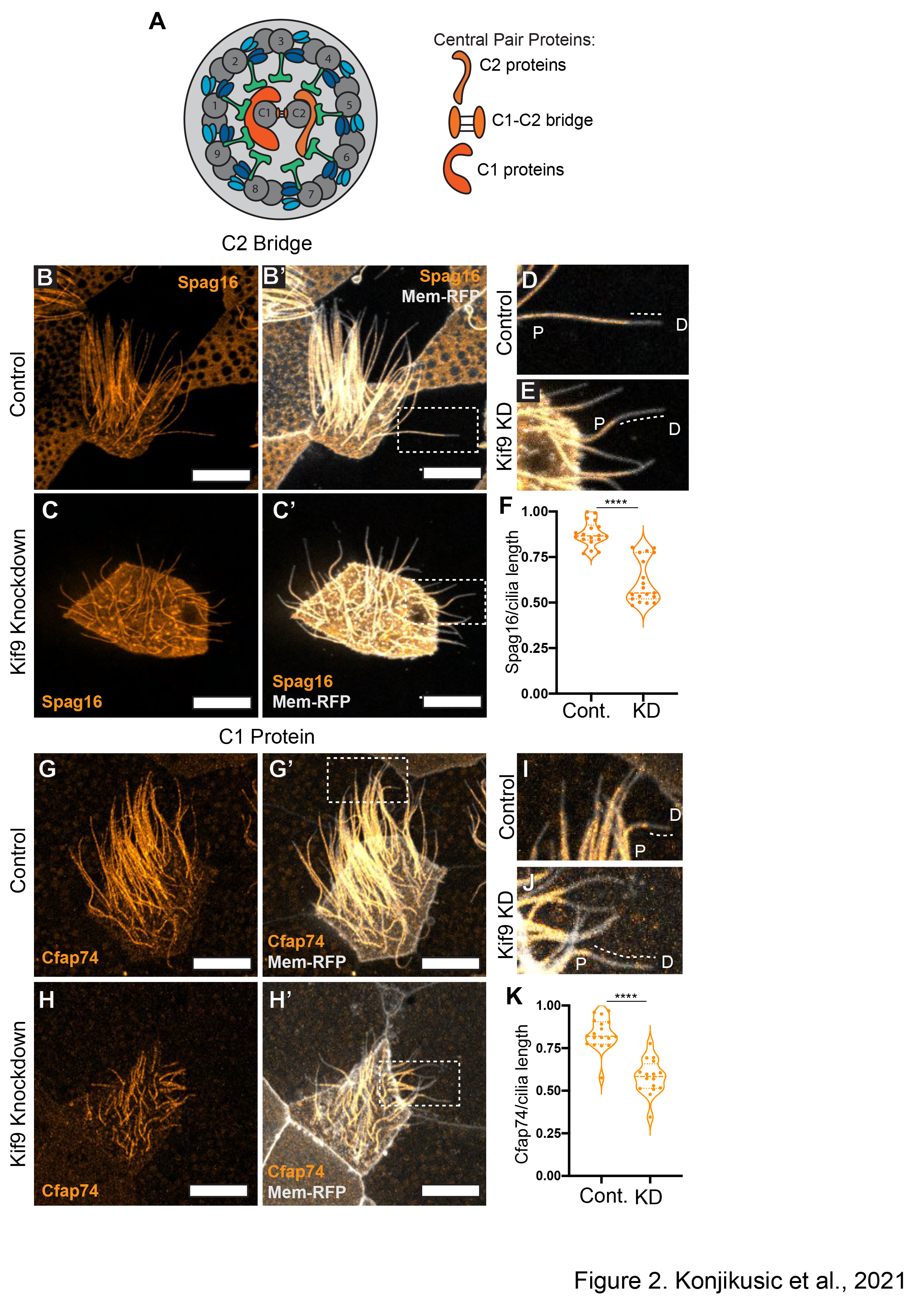
Kif9 contributes to central pair proteins and distal tip compartmentalization. A. Schematic of 9+2 motile ciliary structure with central pair components schematized by tubule association. B-B’. Confocal images of GFP-Spag16 (orange) and membrane-RFP (grays) localization in *Xenopus* Multiciliated cells in Control (B-B’), Kif9 Morpholino (C-C’) injected embryos. D-E. Magnified insets of distal tips of cilia in Control (D) and Kif9 Knockdown (E) cilia showing loss of GFP-Spag16 (orange) from the distal tip of cilia. Dotted line outlining lack of protein at distal ends. P = Proximal, D = Distal. F. Quantification of GFP-Spag16 length over total ciliary length in Control and Kif9 Knockdown injected embryos. P-value = **** < 0.0001. G-H’. Confocal images of GFP-Cfap74 localization in Control (G-G’) and Kif9 Morpholino (H-H’) injected embryos. I-J. Magnified insets of distal tips of cilia in Control (I) and Kif9 knockdown (J) cilia showing loss of GFP-Cfap74 (orange) from the distal tip of cilia. Dotted line outlining lack of protein at distal ends. P = Proximal, D = Distal. K. Quantification of GFP-Cfap74 length over total ciliary length in Control and Kif9 knockdown embryos. All scale bars 10 μm.

To quantify this loss, we measured the length of the cilium containing Spag16 protein and normalized it to total ciliary length as indicated by membrane-RFP, which allowed us to calculate the proportion of the cilium occupied by Spag16 and adjust for ciliary length differences (Figure 2F, Supp. Figure 2). The average proportion Spag16 occupied along the cilium in controls was 87.7%, while Kif9 KD reduced this value to 61.5% of axoneme length (P-value < 0.0001). Additionally, we quantified fluorescence intensity along the axonemes and observed that there was little difference in fluorescence intensity between Kif9 KD and controls in regions where Spag16 remained (Supp. Figure 2B).

We next examined the C1-associated protein Cfap74, variations in which can cause human Primary Ciliary Dyskinesia (PCD) (Dai et al., 2020; Zhao et al., 2019; Sha et al., 2020). Kif9 KD elicited a defect in Cfap74 localization very similar to that observed for Spag16, with loss specifically from the distal axoneme (Figure 2G-K, Supp. Figure 2C-D). These data suggest that loss of Kif9 leads to defects in CA protein localization at distal ends of motile cilia.

### Loss of Kif9 affects the proximal to distal patterning of Radial Spokes, Inner Dynein Arms and Outer Dynein Arms in the cilium

Recent proteomic experiments with *Chlamydomonas* central pair mutants identified a close physical association between Kif9 and a radial spoke “head” protein, Rsph10B/Cfap266 (Dai et al., 2020; Zhao et al., 2019; Yang et al., 2006). We therefore asked if Rsph10B localization may be dependent on Kif9 in *Xenopus* multiciliated cells. We co-injected GFP-Rsph10B and membrane-RFP into control and Kif9 morphant embryos. Strikingly, we observed a reduction in the levels of Rsph10B at distal ends of cilia (Figure 3A-E, Supp. Figure 2E-F, P-value = 0.003), similar to that of Spag16 and Cfap74 (Figure 2). However, in contrast to the central pair proteins, we also observed an increase in fluorescence intensity of Rsph10B in the proximal axoneme of Kif9 morphants when compared to controls (Supp. Figure 2F).

**Figure 3.**
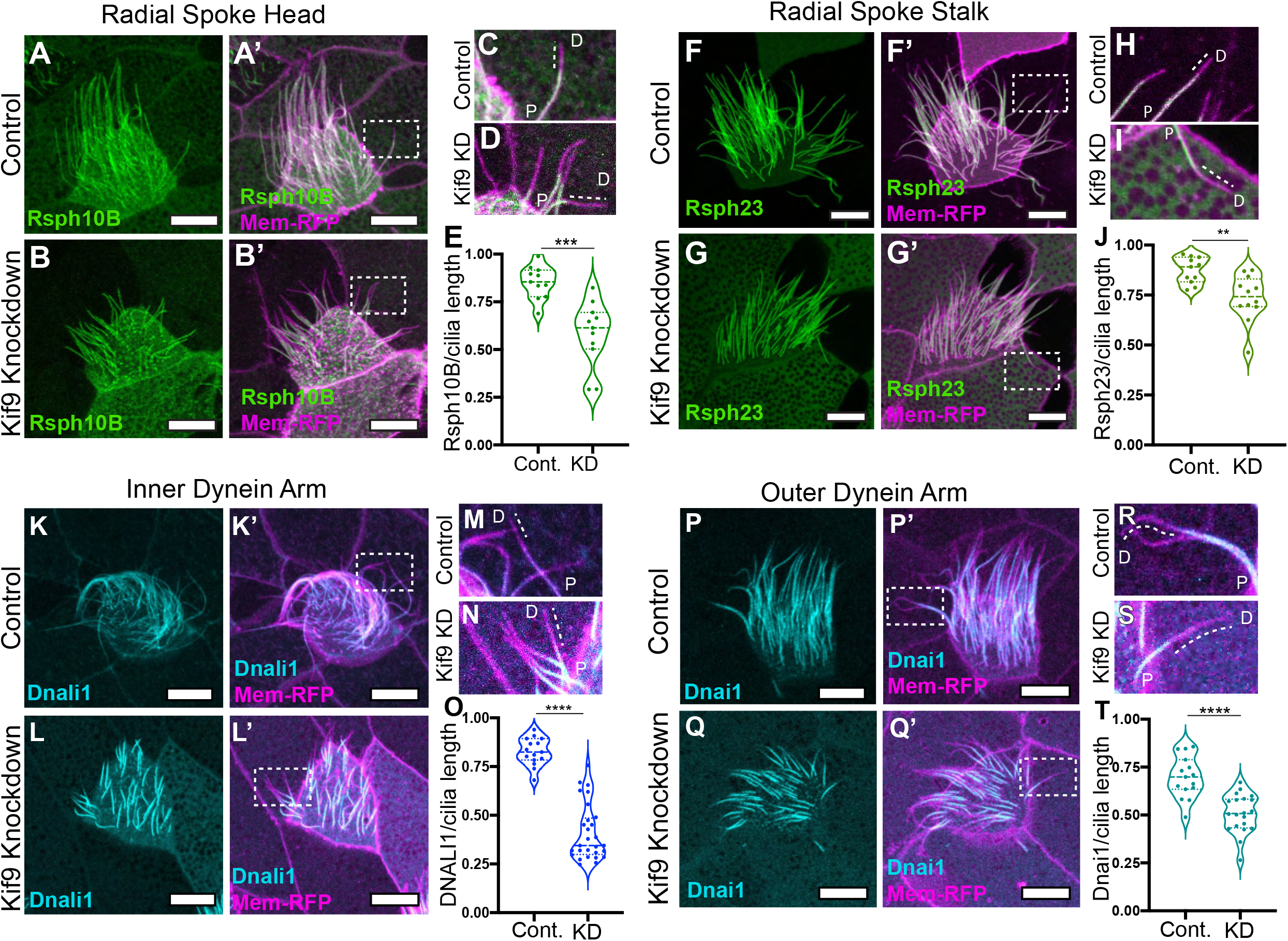
Kif9 is necessary for Radial Spoke, Inner Dynein Arm and Outer Dynein Arm placement in the distal ends of cilia. A-B’. Confocal images of radial spoke head component GFP-Rsph10B (green) localization in Control (A-A’) and Kif9 Morpholino (B-B’) injected embryos. C-D. Magnified insets of cilia in Control (C) and (D) Kif9 knockdown cilia showing loss of GFP-Rsph10B (green) from the distal ends of cilia. Dotted line outlining lack of protein at distal ends. P = Proximal, D = Distal. E. Quantification of GFP-Rsph10B length over total ciliary length in Control and Kif9 knockdown embryos. F-G’. Confocal images of radial spoke stalk component GFP-Rsph23 (green) localization in Control (F-F’) and Kif9 Morpholino (G-G’) injected embryos. H-I. Magnified insets of cilia in Control (H) and Kif9 knockdown (I) cilia showing loss of GFP-Rsph23 from the distal ends of cilia. Dotted line outlining lack of protein at distal ends. P = Proximal, D = Distal. J. Quantification of GFP-Rsph23 length over total ciliary length in Control and Kif9 knockdown embryos. K-L’. Confocal images of inner dynein arm Dnali1-GFP localization in Control (K-K’) and Kif9 Morpholino (L-L’) injected embryos. M-N. Magnified insets of cilia in Control (M) and Kif9 knockdown (M) cilia showing loss of Dnali1-GFP from the distal ends of cilia. Dotted line outlining lack of protein at distal ends. P = Proximal, D = Distal. O. Quantification of GFP-Dnai1 length over total ciliary length in Control and Kif9 knockdown embryos. P-Q’. Confocal images of outer dynein arm GFP-Dnai1 (cyan) localization in Control (P-P’) and Kif9 Morpholino (Q-Q’) injected embryos. R-S. Magnified insets of cilia in Control (R) and Kif9 knockdown (S) cilia showing loss of GFP-Dnai1 (cyan) from the distal ends of cilia. Dotted line outlining lack of protein at distal ends. P = Proximal, D = Distal. T. Quantification of GFP-Dnai1 length over total ciliary length in Control and Kif9 knockdown embryos. All scale bars 10uM.

To ask how loss of Kif9 may affect other members of the radial spoke complex, we tested Rsph23, a radial spoke stalk or “neck” protein that is out of direct contact from the central pair apparatus (Frommer et al., 2015). Live confocal imaging of control and Kif9 KD embryos revealed a striking reduction in Rsph23 along the axonemes (Figure 3F-G’). This reduction in the axoneme stemmed from a loss of Rsph23 from the distal ends of the axonemes (Figure 3H-J, P-value = 0.0023, Supp. Figure 2G-H). Interestingly, this decrease of Rsph23 placement along the axoneme was less prominent than the loss of Rsph10B or any of the central pair proteins we tested (Figure 2, Figure 3A-E).

Radial Spoke proteins function to control dynein driven beating, and most radial spoke mutants in many species displayed paralyzed flagella (Viswanadha et al., 2017). We therefore asked if Kif9 had any effect on inner and outer dynein arm localization in the axoneme by examining GFP-Dnali1 or GFP-Dnai1, respectively. Strikingly, we observed that both inner dynein arm Dnali1 and outer dynein arm Dnai1 were reduced in distal axonemes (Figure 3K-T).

### Kif9 contributes to distal tip integrity in motile axonemes

The specific loss of beating machinery from the distal region of motile cilia led us to ask if Kif9 may contribute to the specialized structures of the axoneme distal tip (Soares et al., 2019). We therefore examined the localization of Spef1, a microtubule bundling protein that is both enriched in the distal tips of *Xenopus* multiciliated cilia and is required for central pair assembly in mammals (Gray et al., 2009; Chan et al., 2005; Zheng et al., 2019). First, we found that unlike other beating machinery, Kif9 protein extended distally in the region marked by enriched Spef1. Unlike Spef1, however, Kif9 was not enriched in this region but rather displayed a mosaic, speckled pattern that was not consistent from cilium to cilium (Figure 4B-C”).

**Figure 4.**
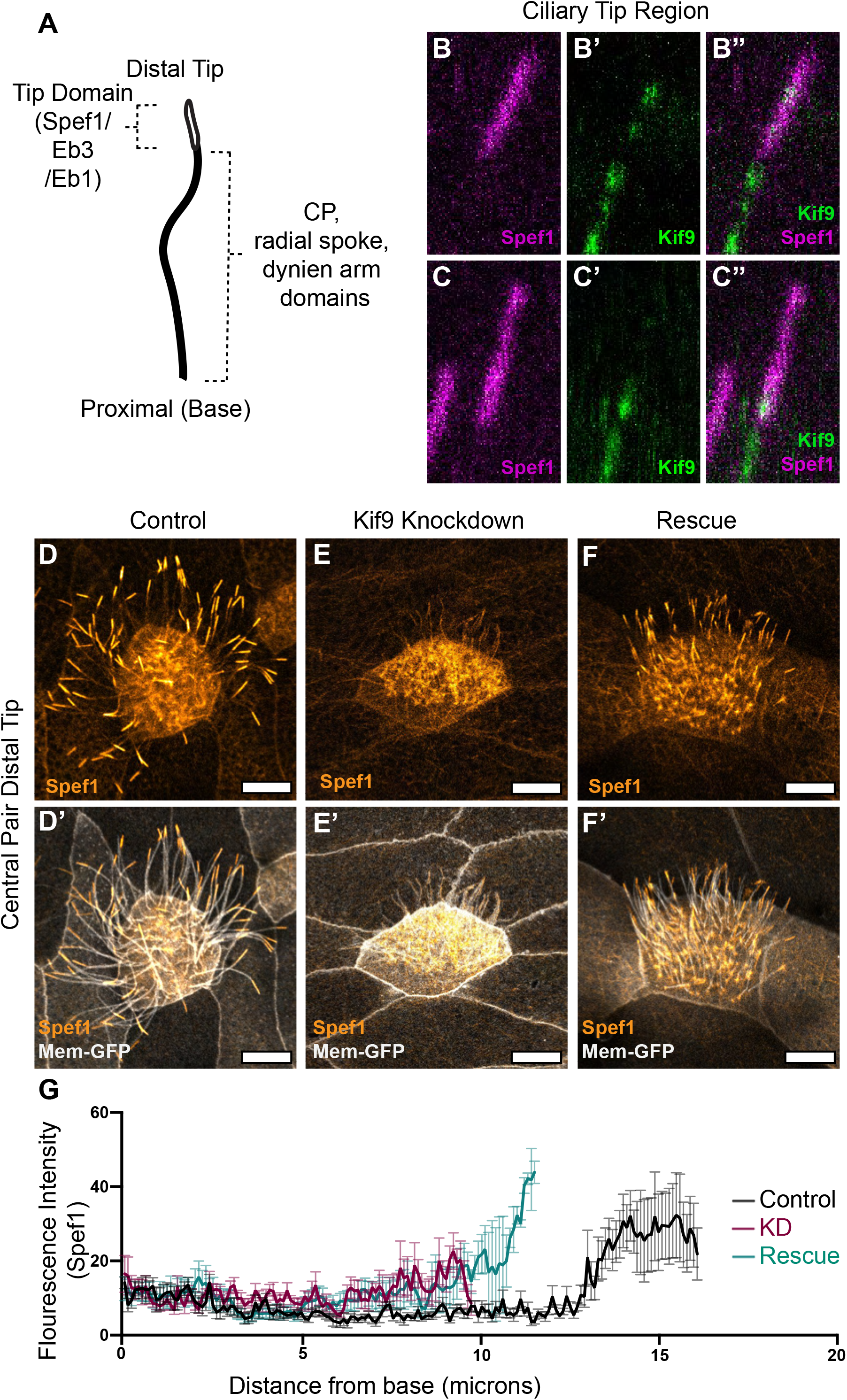
Kif9 contributes to distal tip integrity. A. Schematic of proximal to distal patterning of the motile cilium. Tip domain schematized with proteins known to localize to that domain, as well as proteins localizing to the rest of the axonemes. B-C”. Confocal Images of RFP-Spef1 (magenta) and Kif9-GFP (green) colocalization in the distal tips of the cilium. D-F’. Confocal images of RFP-Spef1 (orange) and Membrane-GFP (grays) in Control (D-D’), Kif9 morpholino injected (E-E’), and Rescue (F-F’) embryos. G. Graph of fluorescence intensity line traces of RFP-Spef1 in Control, Knockdown, and Rescue cilia. All scale bars 10 microns.

More importantly, by co-expressing RFP-Spef1 with membrane-GFP we observed a striking loss of the Spef1-enriched tip domain in Kif9 knockdown cilia (Figure 4D-E’). This phenotype was specific, as it was recused by re-introduction of Kif9 mRNA (Figure 4F-F’). We quantified this loss using fluorescence intensity traces along the cilium (Figure 4G).

Finally, this phenotype did not reflect a general defect in the distal cilium tip, as other proteins that decorate this region were unaffected after Kif9 loss. For example, the microtubule plus-end binding protein Eb3 displays a broad distal enrichment similar to Spef1 (Fig. 5), while Eb1 displays a more concentrated enrichment at the extreme distal end of the cilium (Fig. 5). Both Eb1 and Eb3 remained distally enriched after Kif9 loss. These data argue that Kif9 contributes to the localization of specific elements of the distal tip of motile axonemes.

**Figure 5.**
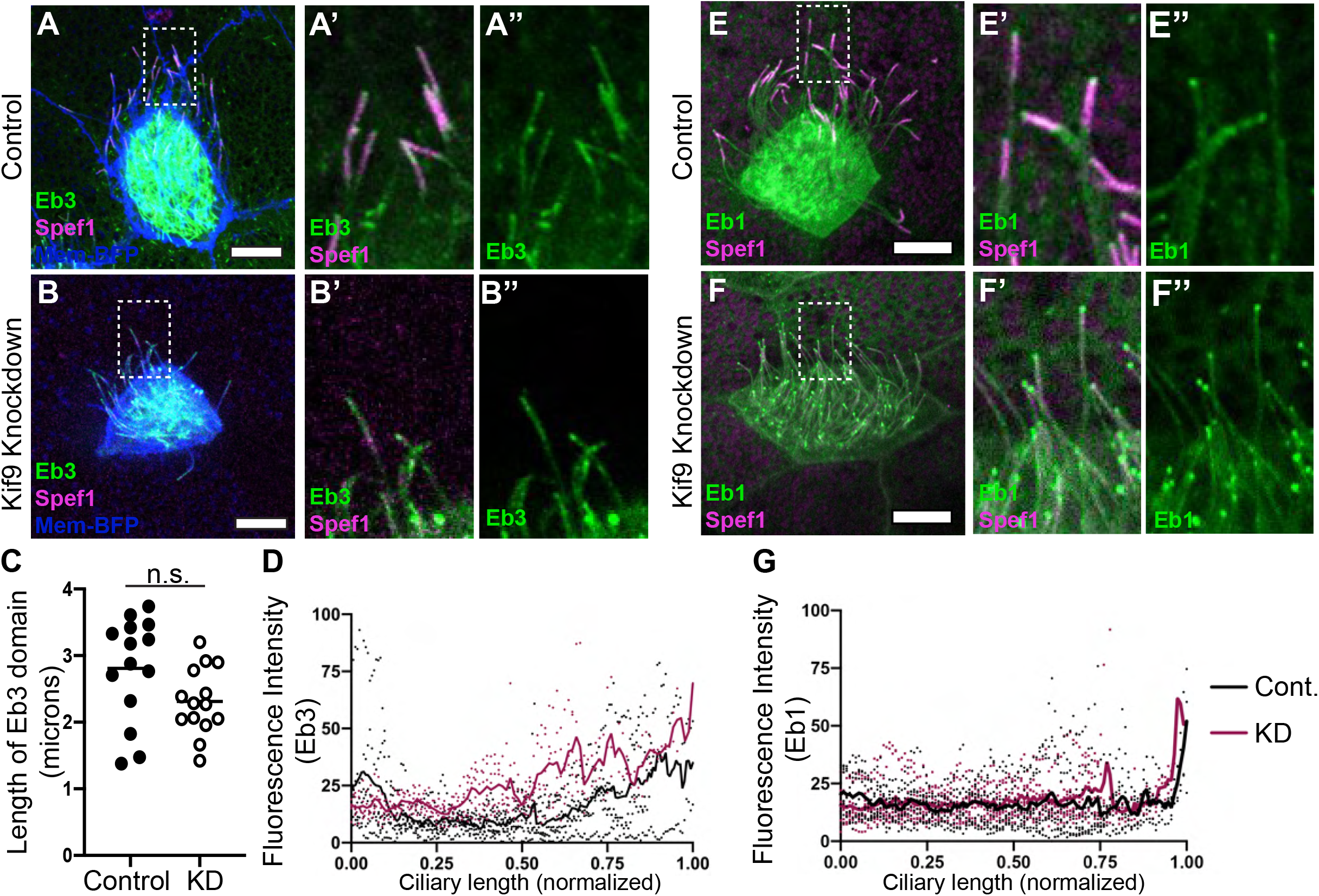
Microtubule plus-end binding proteins Eb1 and Eb3 remain unaffected in the distal tip by knockdown of Kif9. A-B”. Confocal images of Control (A-A”) and Kif9 morpholino injected embryos (B-B”) with Eb3-GFP (green), RFP-Spef1 (magenta), and Membrane-BFP (blue). A’-B’. Insets of ciliary tips with Eb3-GFP (green) an RFP-Spef1 (magenta) in Control and Knockdown cilia. A”-B’’. Insets with Eb3 alone in Control and Knockdown cilia. C. Length measurements of Eb3 domain in Control and Kif9 knockdown embryos. P-value = 0.056. D. Graph of line traces of fluorescence intensities of Eb3 in Control and Knockdown cilia. E-F”. Confocal images of Control (E-E”) and Kif9 morpholino injected embryos (F-F”) with Eb1-GFP (green), RFP-Spef1 (magenta), and Membrane-BFP (blue). E’-F’. Insets of ciliary tips with Eb1-GFP (green) an RFP-Spef1 (magenta) in Control and Knockdown cilia. E”-F”. Insets of Eb1 alone in Control and Knockdown cilia. G. Graph of line traces of fluorescence intensities of Eb1 in Control and Knockdown cilia. All scale bars 10 microns.

### Kif9 can bind microtubules and has slow processivity *in vitro* but does not display processivity in axonemes

Kinesins play diverse cellular roles. While most kinesins acts as motors for intracellular trafficking, others function as microtubule associated proteins or microtubule depolymerizing proteins (Konjikusic et al., 2021). Klp1 binds microtubules *in vitro* (Yokoyama et al., 2004), but its motor activity and the microtubule interactions of its vertebrate ortholog, Kif9, remain unknown. We therefore turned to single-molecule motility assays to assess Kif9 and its behavior on microtubules. Full length *Xenopus* Kif9 showed no ability to bind microtubules *in vitro* (Figure 6A, B-B”), consistent with the fact that full-length kinesin motors are often autoinhibited (Verhey and Hammond, 2009; Brunnbauer et al., 2010). By contrast, a truncated version of Kif9 containing only the motor domain and a portion of the coiled-coiled domain bound strongly to microtubules (Figure 6A’, 6C-C”). Moreover, live time-lapse imaging revealed processive movement of Kif9 along microtubules in a unidirectional manner (Movie 7, Figure 6D-D” arrow following Kif9 puncta). Kymographs confirmed processivity *in vitro*, albeit at relatively slow rates (∼8nm /sec; Figure 2E, Supp. Figure 3).

**Figure 6.**
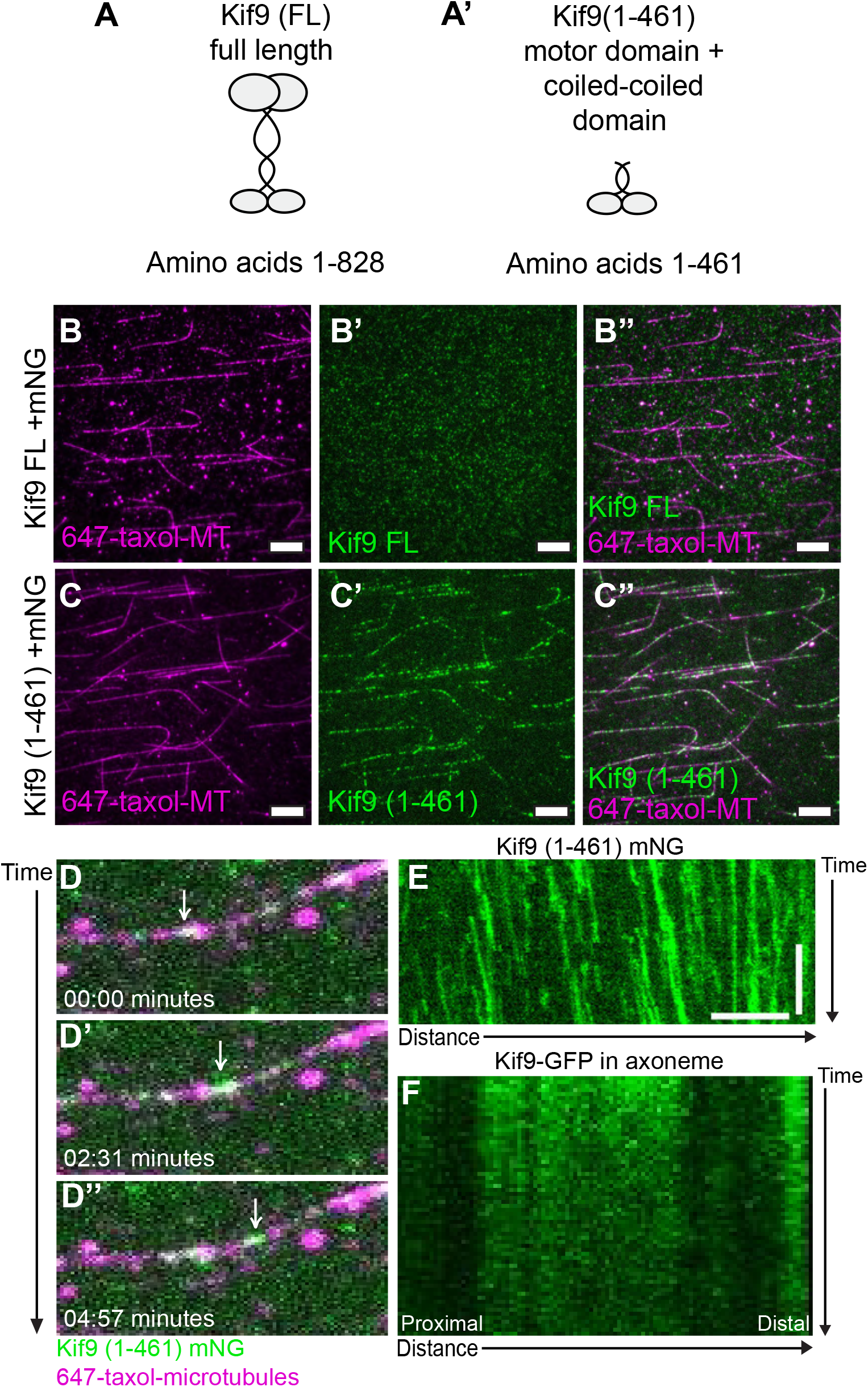
Motor activity of *Xenopus* Kif9. A. Schematic of *Xenopus laevis* Kif9 constructs made for single molecule assays. B-B’’. Images of single molecule microtubule binding assays for full length Kif9 (Kif9 (FL) (green)) on taxol stabilized microtubules (magenta). All scale bars 10 μm. C-C’’. Images of single molecule microtubule binding assays for a truncated version (Kif9 (1-461)) (green) on taxol stabilized microtubules (magenta). All scale bars 10 μm. D-D’’. Still frames from live imaging of Kif9 processivity on microtubules in minutes. White arrow follows Kif9 particle (green) movement along microtubule (magenta). E. Kymograph of live imaging of Kif9 (1-461)-mNG *in vitro* on microtubules. Scale bars 2 minutes and 5 μm. F. Kymograph of live imaging of Kif9-GFP construct *in vivo* in *Xenopus laevis* multiciliated cell axonemes.

These results suggest that Kif9 could act as a traditional transport motor to carry ciliary components along the central pair microtubules. To test this, we expressed Kif9-GFP in motile axonemes and performed time-lapse imaging. However, at mature multiciliated cell stages, we observed no processivity of Kif9 *in vivo*, as seen in both movies and kymographs (Movie 8, Figure 6F). Thus, our data show that although Kif9’s motor domain is capable of processive motility, the protein does not undergo processive motility along axonemal microtubules.

## Discussion

Ciliary beating requires the action of the inner and outer dynein arms, the radial spokes, and the central pair, but how these systems work in concert remains poorly understood. Here, we extend previous studies of the central pair kinesin Klp1 in *Chlamydomonas* by showing that the vertebrate ortholog Kif9 is not only required for ciliary beating, but also has processive motor activity and is required for the organization of the distal axoneme in multiciliated cells. The work provides several new insights into the cell biology of ciliary beating.

First, the function of the central pair has been postulated to control beat and waveform through mechanical links to radial spoke and inner and outer dynein arms (Oda et al., 2014). Several studies have shown a physical link between central pair apparatus and radial spoke head proteins that occurs when cilia are undergoing “bending” during beat (Goodenough and Heuser, 1985; Oda et al., 2014; Warner, 1970). However, which proteins create this link is unknown. One hypothesis is that a microtubule motor protein, such as a kinesin motor, could serve this function as it could bind the central pair of microtubules and adjacent proteins and use its motor activity to fine tune beat and waveform through motor action, much like the action of axonemal dyneins (Viswanadha et al., 2017). Interestingly, our single-molecule analysis shows that Kif9 displays slow processivity *in vitro*, arguing that it may function as a conventional motor for intracellular trafficking. Yet, in mature multiciliated cells we observed no processivity in axonemes. This result is similar to that of axonemal dynein, which have processivity *in vitro* but participate strictly in force generation that regulates ciliary beating *in vivo* (Sakakibara et al., 1999). This leads us to hypothesize that instead of functioning as a conventional motor for axonemal transport, Kif9/Klp1 may contribute to ciliary beating and waveform through a similar mechanism as dynein arms generating force through motor action.

In support of this hypothesis, proteomic studies suggest a close association of Kif9 with the central pair protein Pf20/Spag16 as well as the radial spoke protein Fap266/Rsph10B (Dai et al., 2020; Zhao et al., 2019). The observed loss of both of these proteins from the distal axoneme after Kif9 knockdown support this idea (Figure 7B). Altogether, we hypothesize that Kif9 may act as a physical link between the central pair and the radial spoke, fine tuning waveform and beat frequency through its motor domain, while interacting with radial spoke to orchestrate beating.

**Figure 7.**
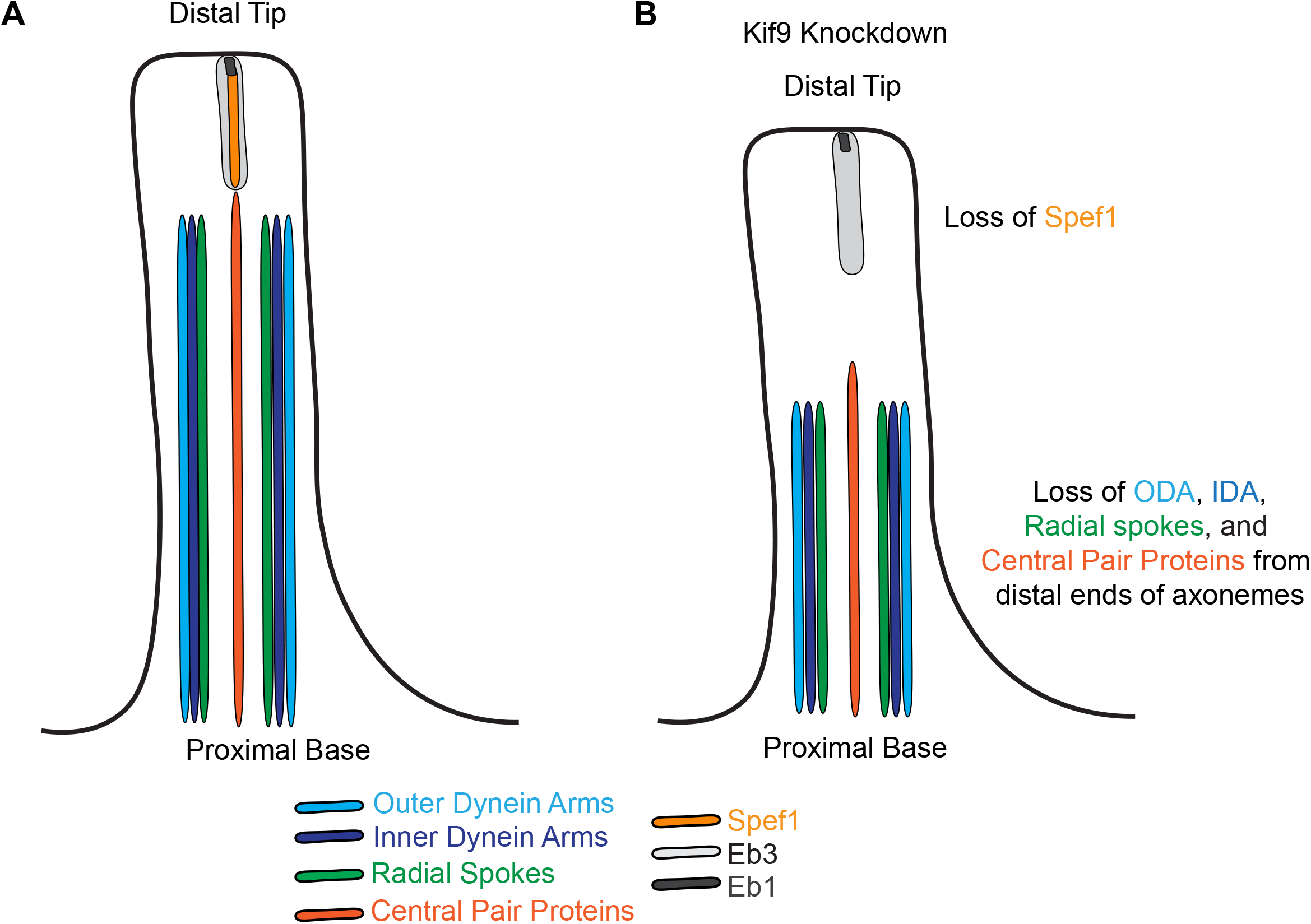
Summary of effect of Kif9 knockdown on proximal to distal patterning of the motile cilium. A. Schematic of wildtype motile cilia in *Xenopus* with domains schematized based off findings. B. Schematic of Kif9 knockdown cilia and changes to domains observed. C.? Legend of colors that correlate to specific domains and proteins within the schematic for A and B.

Motile axonemes also display proximodistal patterning, including a specific tip domain (Figure 7A, (Soares et al., 2019)). Here, we further define the specific domains of dynein arms, radial spokes, and CA proteins in *Xenopus* multiciliated cells, and also provide a more detailed understanding of the distal axoneme tip (Figure 7A). Several different types of “capping” structures have been described in mammalian and *Xenopus* multiciliated cells (Soares et al., 2019; Dentler and Lecluyse, 1982; Portman et al., 1987; LeCluyse and Dentler, 1984). What proteins contribute to these structures remains unclear and varies among tissue and cilia types (Soares et al., 2019). Spef1 localizes to *Xenopus* multiciliated cells in a enriched domain at the distal tips of cilia ((Gray et al., 2009), Figure 4A), and later experiments in mouse found that Spef1 is necessary for central pair formation in mouse testes and ependymal cell culture and functions in vitro as a microtubule bundling protein (Chan et al., 2005; Zheng et al., 2019, 1).

Our new work reveals that loss of Kif9 leads to loss of the Spef1 distal tip domain in *Xenopus* multiciliated cells, though two other distal tip proteins, Eb1 and Eb3 remain unaffected (Figure 7B). We suggest that Spef1 may be a central pair distal tip contributor in multiciliated cells, and that proper CA assembly may be necessary for tip integrity in multiciliated cells. Interestingly, several different central pair mutants in *Chlamydomonas* have shorter cilia (Lechtreck et al., 2013). We speculate this decrease in length may come from loss of distal tip integrity in these mutants.

In conclusion, we show that is both a processive motor and is required for the normal localization of three major motility complexes in motile axonemes. Our argue that the CA may be necessary hierarchically for placement of radial spoke and inner and outer dynein arms into the axonemes.

## Methods and Materials

### *Xenopus* embryo manipulations

*Xenopus* embryo manipulations were carried out using the standard protocols. Briefly, female adult *Xenopus* were induced to ovulate by injection of hCG (human chorionic gonadotropin). *In vitro* fertilization was carried out by homogenizing a small fraction of a testis in 1/3 X Marc’s Modified Ringer’s (MMR). Embryos were de-jellied in 1/3x MMR with 2.5-3% L-cysteine (pH 7.8-8). Embryos were microinjected with mRNA and morpholinos in 2% Ficoll (w/v) in 1/3x MMR. Embryos were washed with 1/3x MMR 1 hour post injection. Each injection was repeated 3-4 times across several days and clutches of embryos.

### Microinjections

Gene sequences were obtained from Xenbase (http://www.xenbase.org). Open reading frames (ORF) of genes were amplified from the *Xenopus* cDNA library by polymerase chain reaction (PCR), and then inserted into a pCS vectors containing a fluorescence tag. The following constructs were cloned into pCS10R vector: GFP-dnai1, GFP-nme5 (Rsph23), RFP-clamp (spef1), GFP-spag16, GFP-Cfap74, GFP-Rsph10b, GFP-dnali1, eb3-GFP from (Shindo et al., 2008), eb1-GFP, membrane-RFP, membrane-GFP, membrane-BFP, Centrin4-RFP, and kif9-GFP. These constructs were linearized and capped mRNAs were synthesized using mMESSAGE/mMACHINE SP6 transcription kit. Between 50-100pg of any given mRNA was injected into the two ventral blastomeres at the 4 cell stage. For rescue experiments 600pg of kif9 mRNA was injected. The kif9 morpholino was designed as a splice blocking morpholino to exon 1 of the kif9 mRNA sequence. The working concentration was 15ng and sequence as follows: 5’ ATTCTCATATTCAATAGTCTTACCT 3’.

### Imaging and Analysis

Xenopus embryos were grown up to stage 22-25 and mounted onto an imaging chamber in 1/3 MMR and live imaged immediately. The Bead flow movies and live cilia beat videos were taken on a Nikon W1 Spinning Disc Confocal at an image rate of 12.5 ms per frame. Live still images of axonemes were captured with a Zeiss LSM700 laser scanning confocal microscope with a 63x oil objective. Each experiment was repeated 3-4 times across several days and clutches of embryos. Approximately 15 different images (3-5 embryos) were taken for quantification, 5 different cilia per cell and at least three cells per image were quantified.

Beads (or microspheres) of size 1 um are added to the fluid medium for the purpose of tracing/visualizing and quantifying the fluid flow induced by the beating cilia. Bead flow movies were subjected to Flowtrace (Gilpin et al., 2017), for visualization of bead flow patterns across the epidermis. For this Flowtrace analysis, we selected an interrogation window size of 50 frames.

To quantify the fluid flow speeds, we carried out a Particle Image Velocimetry (PIV) analysis using the Matlab-based PIVlab package(Thielicke and Stamhuis, 2014). First, we extracted individual frames from the time-lapse datasets (movies) and we carried out the image preprocessing step by selecting the contrast limited adaptive histogram equalization (CLAHE) option (window size 20 px). Next, we proceeded with the PIV analysis using these settings: FFT window deformation, Interrogation area 128 × 128 pixels with 50% overlap (Pass 1), and with interrogation area of 64 × 64 pixels with 50% overlap (Pass 2). After this step, the vector validation was done by choosing suitable velocity limits, standard deviation filters and interpolation of missing data. We carefully selected a rectangular region just adjacent and parallel to the ciliary beating surface on the embryo. We quantified the average speed in this selected region over time, and the error bars represent the standard deviation. This procedure was utilized for each individual dataset for the control, Kif9 and rescue cases. To obtain a characteristic speed for each case (control, Kif9 and rescue), further averaging was carried out: for control and Kif9 (N = 5 datasets), and for rescue (N = 10 datasets).

Quantification of cilia length, length of each construct along axonemes, and fluorescence intensity profiles were all taken in FIJI. Kymographs of cilia beat were also generated using FIJI. Briefly, lines were drawn, taking length measurements and fluorescence intensity profiles. Outputs were placed into GraphPad Prism 8 and graphs were generated. All statistical analyses were done in GraphPad Prism 8.

### *Xenopus* immunostaining

For immunostaining, protocols previously described in (Brooks and Wallingford, 2015) were followed. Briefly, wildtype and MO injected embryos were fixed with 4% PFA at stages 22-25 for 1 hour at RT. They were then washed with PBS + 0.1% Tween (PBST), and subsequently washed in MeOH. They were dehydrated and permeabilized in 100% MeOH at -20 degrees C for 30 minutes to 1 hour. Embryos were then rehydrated and washed with PBST. Embryos were then blocked for 1 hr at RT in 10% NGS, 5% DMSO in 1x PBST. Primary antibodies were incubated overnight and embryos were washed 3x in PBST, then incubated with secondary antibody for 1 hour at RT. A rabbit Kif9 antibody was used at 1:100 dilution from Atlas Antibodies (HPA022033). Mouse acetylated tubulin antibody was incubated at 1:1000 from Sigma (6-11B-1). Secondaries Alexa fluorophores anti-rabbit 488 and anti-mouse 647 were used at 1:1000 dilutions.

### RT-PCR

To verify the efficiency of our *Kif9* Morpholino, MO was injected into all 4 cells at the 4 cell stage and total RNA was isolated using the TRIreagent (Thermofischer) at stage 23. One microgram of mRNA was used to synthesize cDNA using iScript (Biorad). Kif9 and B-actin cDNAs were amplified by GoTag Green Master Mix (Promega) and with the following primers:

Bactin F1: 5’ GCCCGCATAGAAAGGAGACAG 3’

Bactin F2: 5’ CCAAACCTCGCTCAGTGACC 3’

Bactin R1: 5’ TCATCCCAGTTGGTGACAATGC 3’

Bactin R2: 5’ TCCCATTCCAACCATGACACC 3’

Kif9 F1: 5’ GAGACGGGATAGTTTACACACAGC 3’

Kif9 R1: 5’ TGGAGGCAAGGTTTAGGGATAAGC 3’

Kif9 F2: 5’ CGCTGAAGCCAAGAGCTGAAC 3’

Kif9 R2: 5’ GCATCTGGAACAGTGGAAAGGAG 3’

### Airway epithelial cell culture and immunostaining

Human airway tracheas were retrieved from surgical excess of tracheobronchial segments of lungs donated for transplantation. Human tracheobronchial epithelial cells (hTEC) were isolated from sections of normal human trachea obtained from non-smoking donors lacking respiratory pathology. These unidentified tissues are exempt from regulation by HHS regulation 45 CFR Part 46. hTEC cells were expanded in-vitro and allowed to differentiate into ciliated cells using air-liquid interface (ALI) conditions on supported membranes (Transwell, Corning Inc., Corning, NY), as previously described (PMID: 12388377). Paraffin embedded tracheal sections or cultured primary airway cells were fixed and immunostained as previously described (PMID: 17488776). Nuclei were stained using 4’, 6-diamidino-2-phenylindole (DAPI, Vector Laboratories, Burlingame, CA, USA). KIF9, was detected using primary antibodies obtained from MilliporeSigma (St. Louis, MO, HPA022033). Basal bodies were detected using antibodies against Centrin (clone 20H5, MilliporeSigma). DNAI1, a marker of outer dynein arm, was detected using NeuroMab (UC Davis, Ca, clone UNC 65.56.18.11). Images were acquired using an epifluorescent microscope interfaced with imaging software (LAS X; Leica) and adjusted globally for brightness and contrast using Affinity Photo (Serif Ltd, Nottinghamshire, United Kingdom).

### Single Molecule Microtubule Assays

#### Plasmid

To generate pMT-*xl*KIF9(1-461)-mNeonGreen plasmid, the coiled-coil prediction of full-length *xl*KIF9 was carried out in Marcoil (Delorenzi and Speed, 2002) and COILS (Lupas, 1996). *xl*KIF9(−461) truncation which includes the first two coiled coil domains and is expected to form a dimer was amplified by PCR from full-length *xl*KIF9 and subcloned into pMT-mNeonGreen vector by NEBuilder HiFi DNA assembly cloning kit. The plasmid was verified by DNA sequencing.

#### Cell culture, transfection and lysate preparation

*Drosophila* S2 cells were cultured in Schneider’s *Drosophila* medium (Gibco) supplemented with 10% (vol/vol) FBS (HyClone) at 26°C. Plasmid for expression of *xl*KIF9(1-461)-mNeonGreen in the pMT vector was transfected into S2 cells using Lipofectamine LTX Reagent with PLUS Reagent (Invitrogen) according to manufacturer’s instructions. Expression of *xl*KIF9(1-461)-mNeonGreen was induced by adding 1mM CuSO_4_ to the medium after 4-5h transfection.

To prepare cell lysate for single molecule assays, S2 cells were harvested after 48h transfection. The cells were centrifuged at low speed at 4°C. The cell pellet was washed with PBS buffer and resuspended in ice-cold lysis buffer (25 mM HEPES/KOH, 115 mM potassium acetate, 5 mM sodium acetate, 5 mM MgCl2, 0.5 mM EGTA, and 1% Triton X-100, pH 7.4) freshly supplemented with 1 mM ATP, 1 mM PMSF, 1 mM DTT and protease inhibitors (Sigma-Aldrich). After the cell lysate was clarified by centrifugation at full-speed at 4°C, aliquots of the supernatant were snap frozen in liquid nitrogen and stored at -80°C until further use.

#### Single-molecule motility assays

HiLyte647-labeled microtubules were polymerized from purified tubulin including 10% Hily647-labeled tubulin (Cytoskeleton) in BRB80 buffer (80 mM Pipes/KOH pH 6.8, 1 mM MgCl_2_, and 1 mM EGTA) supplemented with 1 mM GTP and 2.5 mM MgCl_2_ at 37°C for 30 min. 20 μM taxol in prewarmed BRB80 buffer was added and incubated at 37°C for additional 30 min to stabilize microtubules. Microtubules were stored in the dark at room temperature for further use. A flow cell (∼10 μl volume) was assembled by attaching a clean #1.5 coverslip (Fisher Scientific) to a glass slide (Fisher Scientific) with two strips of double-sided tape. Polymerized microtubules were diluted in BRB80 buffer supplemented with 10 μM taxol and then were infused into flow cells and incubated for 5 min at room temperature for nonspecific adsorption to the coverslips. Subsequently, blocking buffer (15 mg/ml BSA and 10 μM taxol in P12 buffer) was infused and incubated for 5min. Finally, kinesin motors in the motility mixture [2 mM ATP, 0.4 mg/ml casein, 6 mg/ml BSA, 10 μM taxol, and oxygen scavenging (1 mM DTT, 1 mM MgCl2, 10 mM glucose, 0.2 mg/ml glucose oxidase, and 0.08 mg/ml catalase) in P12 buffer] was added to the flow cells. The flow-cell was sealed with molten paraffin wax.

Images for single molecule assay were acquired by TIRF microscopy using an inverted microscope Ti-E/B (Nikon) equipped with the perfect focus system (Nikon), a 100× 1.49 NA oil immersion TIRF objective (Nikon), three 20-mW diode lasers (488 nm, 561 nm, and 640 nm) and an electron-multiplying charge-coupled device detector (iXon X3DU897; Andor Technology). Image acquisition was controlled using Nikon Elements software and all assays were performed at room temperature. Images were acquired continuously every 5 s for 5 min. Maximum-intensity projections were generated and the kymographs were produced by drawing along tracks of motors (width= 3 pixels) using Fiji/ImageJ2.

## Figure Legends

**Supplementary Figure 1.**
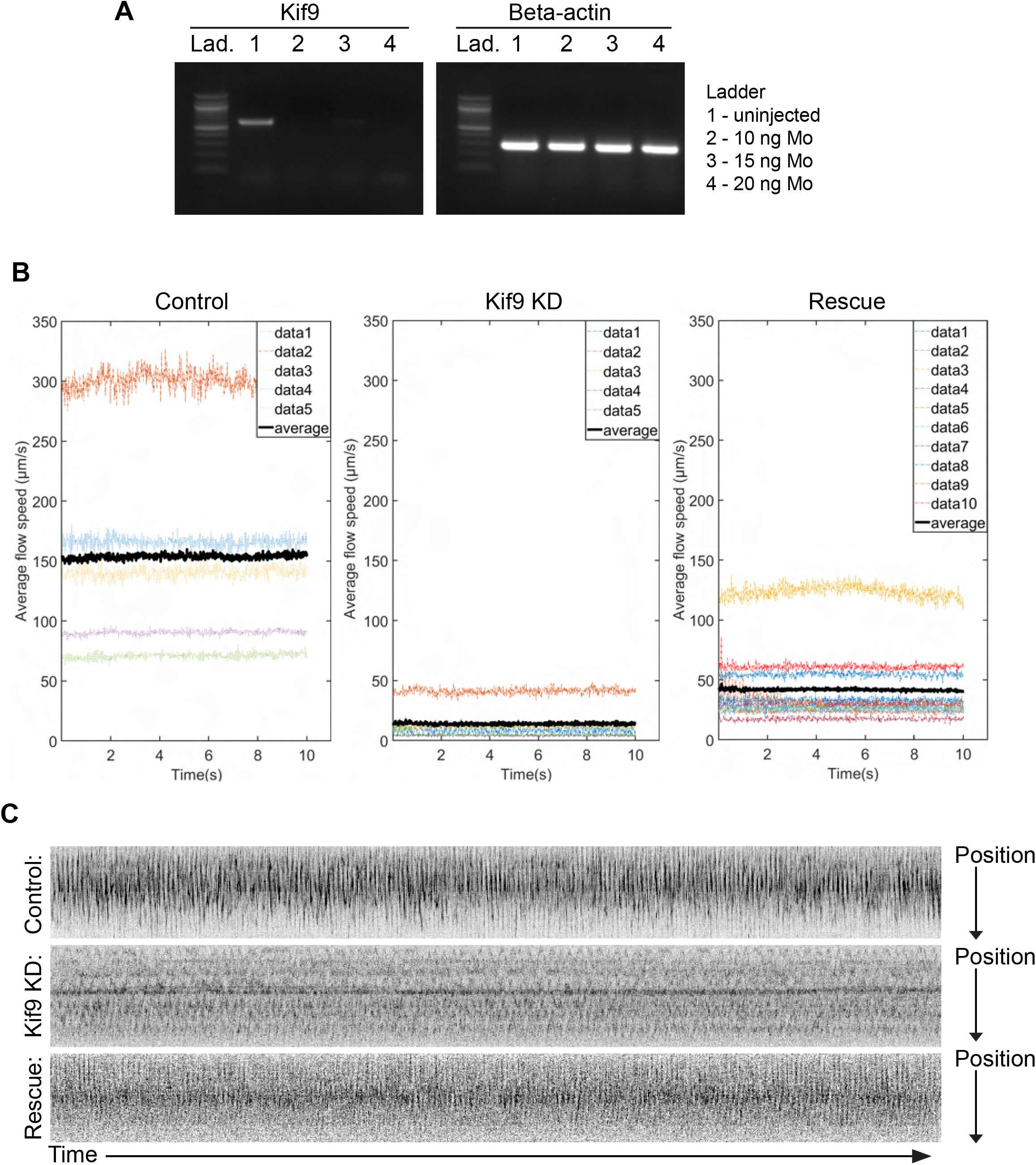
A. Gel image of RT-PCR of Kif9 expression in Kif9 morpholino injected embryos at wildtype, 10 ng, 15 ng, and 20 ng doses. B. Graphs of average bead flow across the epidermis of stage 24 *Xenopus* embryos in Control, Kif9 morpholino, and Rescue injected embryos. C. Kymographs of ciliary beat in Control, Kif9 Morpholino, and Rescue injected constructs.

**Supplementary Figure 2.**
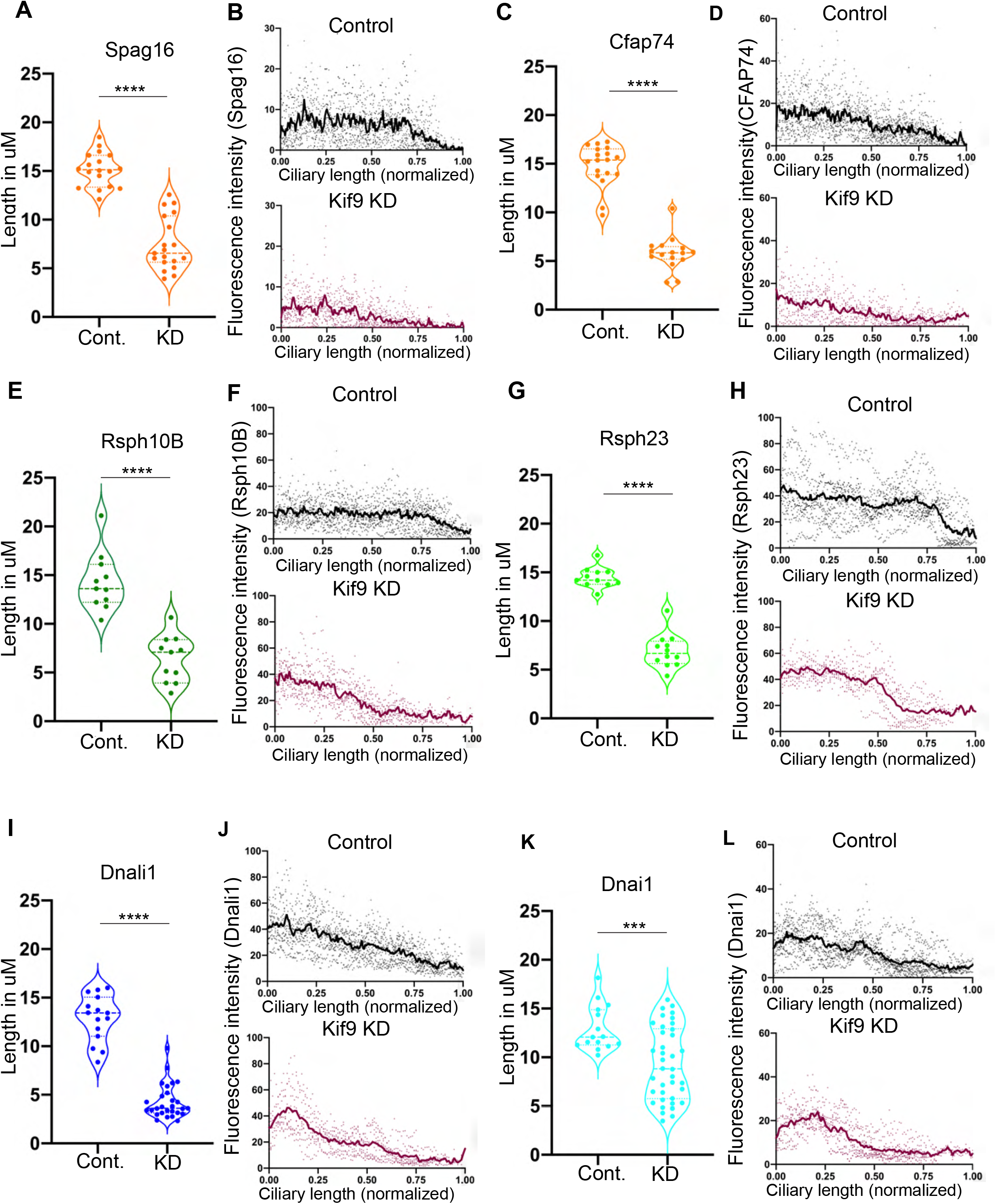
A. Raw quantification of C2 bridge central pair component GFP-Spag16 length in Control and Morpholino injected embryos. B. Graphs of fluorescence intensity of GFP-Spag16 across the length of the cilia, normalized to 1. C. Raw quantification of C1 central pair component GFP-Cfap74 length in Control and Morpholino injected embryos. D. Graphs of fluorescence intensity of GFP-Cfap74 across the length of the cilia, normalized to 1. E. Raw quantification of radial spoke head component GFP-Rsph10B length in Control and Morpholino injected embryos. F. Graphs of fluorescence intensity of GFP-Rsph10B across the length of the cilia, normalized to 1. G. Raw quantification of radial spoke stalk component Rsph23 length in Control and Morpholino injected embryos. H. Graphs of fluorescence intensity of GFP-Rsph23 across the length of the cilia, normalized to 1. I. Raw quantification of inner dynein arm GFP-Dnali1 length in Control and Morpholino injected embryos. J. Graphs of fluorescence intensity of GFP-Dnali1 across the length of the cilia, normalized to 1. K. Raw quantification of outer dynein arm GFP-Dnai1 length in Control and Morpholino injected embryos. L. Graphs of fluorescence intensity of GFP-Dnai1 across the length of the cilia, normalized to 1.

**Supplementary Figure 3.**
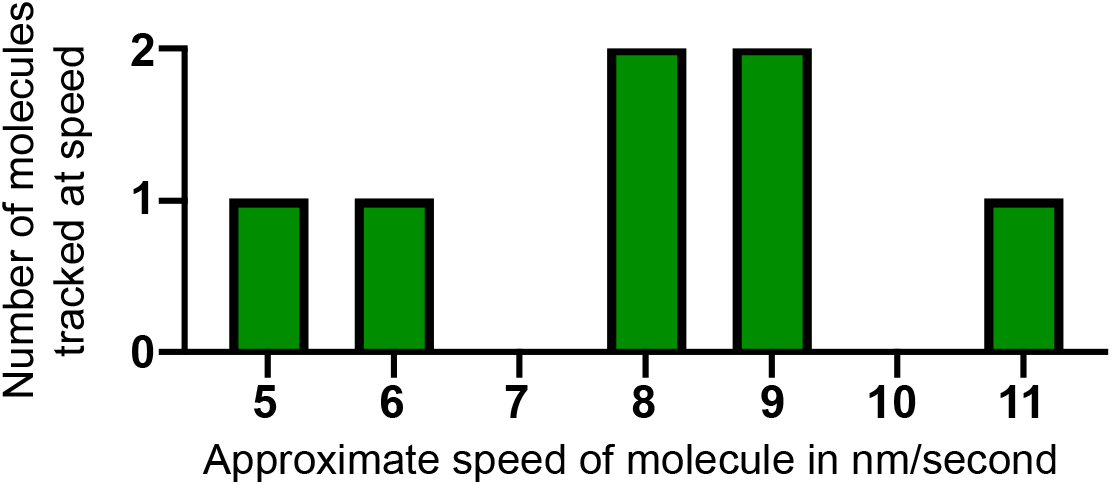
Histogram of approximate speed of truncated Kif9 movement along microtubules *in vitro*. Values binned to 1 nanometer/second. X-axis = number of molecules observed at approximate speed. Y-value = approximate speed of molecule in nm/second.

**Movie 1. Flowtrace movies of beads flowing past epidermis in control embryos**.

**Movie 2. Flowtrace movies of beads flowing past epidermis in Kif9 knockdown embryos.**

**Movie 3. Flowtrace movies of beads flowing past epidermis in rescue treated embryos.**

**Movie 4. Multiciliated cell beating in control embryos**.

**Movie 5. Multiciliated cell beating in Kif9 knockdown embryos.**

**Movie 6. Multiciliated cell beating in rescue treated embryos**.

**Movie 7. Movie of truncated version of Kif9 (Kif9 1-461 mNG) migrating on microtubules.**

**Movie 8. Movie of Kif9-GFP in vivo in *Xenopus* multiciliated cells**.

## Notes

### Competing Interest Statement

The authors have declared no competing interest.

